# Comparative evaluation of *Aspergillus niger* strains for endogenous pectin depolymerization capacity and suitability for D-galacturonic acid production

**DOI:** 10.1101/858548

**Authors:** Dominik Schäfer, Kevin Schmitz, Dirk Weuster-Botz, J. Philipp Benz

**Affiliations:** Technical University of Munich, Department of Mechanical Engineering, Institute of Biochemical Engineering, Boltzmannstr. 15, 85748 Garching, Germany; Technical University of Munich, Holzforschung München, Wood Bioprocesses, TUM School of Life Sciences Weihenstephan, Hans-Carl-von-Carlowitz-Platz 2 85354 Freising, Germany

**Keywords:** *Aspergillus niger*, agricultural residues, sugar beet pulp, pectinase, d-galacturonic acid

## Abstract

Pectinaceous agricultural residues rich in d-galacturonic acid (d-GalA), such as sugar beet pulp, are considered as promising feedstocks for waste-to-value conversions. *Aspergillus niger* is known for its strong pectinolytic activity. However, while specialized strains for production of citric acid or proteins are openly available, this is not the case for the production of pectinases. We therefore systematically compared the pectinolytic capabilities of six *A. niger* strains (ATCC 1015, ATCC 11414, NRRL 3122, CBS 513.88, NRRL 3, N402) using controlled batch cultivations in stirred-tank bioreactors. *A. niger* ATCC 11414 showed the highest polygalacturonase activity, specific protein secretion and a suitable morphology. Furthermore, d-GalA release from sugar beet pulp was 75% higher compared to the standard lab strain *A. niger* N402. Our study therefore presents a robust initial strain selection to guide future process improvement of d-GalA production from agricultural residues and identifies the most suitable base strain for further genetic optimizations.

## Introduction

Global academic and industrial efforts to improve the sustainability of industrial processes for the modern bioeconomy have sparked interest in the utilization of feedstocks that are economically viable, non-food grade and do not compete with food resources [6,22,25,42,47,49,56]. As a result, agricultural waste streams have gained momentum in recent years (reviewed by Amoah [3]). Notably, downstream fermentation products derived from pectin-rich biomass are met with industrial interest by the plastics, cosmetics, and food industry, as recently reviewed by Kuivanen [27] and Schmitz [44].

Apart from harsh thermo-chemical pre-treatment and hydrolysis approaches [12], complex enzyme broths from cultivations of natural pectin-degrading microorganisms can be used for the release of the constituent saccharides, such as d-galacturonic acid (d-GalA) as the main backbone sugar of pectin. The filamentous fungus and saprotroph *Aspergillus niger* (*A. niger*) is a well characterized microorganism for pectin utilization and depolymerization as well as a well-established industrial workhorse with multiple applications in enzyme, citric acid and other organic acid production [9,30]. Pectinases from *A. niger* contribute to a global multi-billion dollar market for biomass-degrading enzymes [9,30,45] with applications ranging from fruit, vegetable and juice processing to textile and paper treatment [21] as well as saccharification for bioethanol production [11,48,54]. However, with today’s perception of d-GalA shifting from an inevitable component of complex biomass feedstocks to a target product for subsequent fermentations, more versatile and higher yielding host strains are needed to extend the range of commercial enzymes, as outlined in a recent white paper on the current challenges of research on filamentous fungi in the context of a sustainable bio-economy [30].

For efficient industrial-scale applications, ideal base strains need to be identified for specific tasks. For *A. niger*, several lineages have been identified and adapted for specific purposes. The most cited lineages of *A. niger* encompass three main clades, namely: (i) strains adapted for easy handling and genetic manipulation in the laboratory environment, which are based on *A. niger* NRRL 3 (CBS 120.49, ATCC 9029); (ii) strains for improved citric acid production, based on *A. niger* ATCC 1015 (NRRL 328, CBS 113.46); and (iii) strains for protein production and secretion, based on *A. niger* NRRL 3122 (CBS 115989; ATCC 22343) (Fig. 1).

**Fig. 1:**
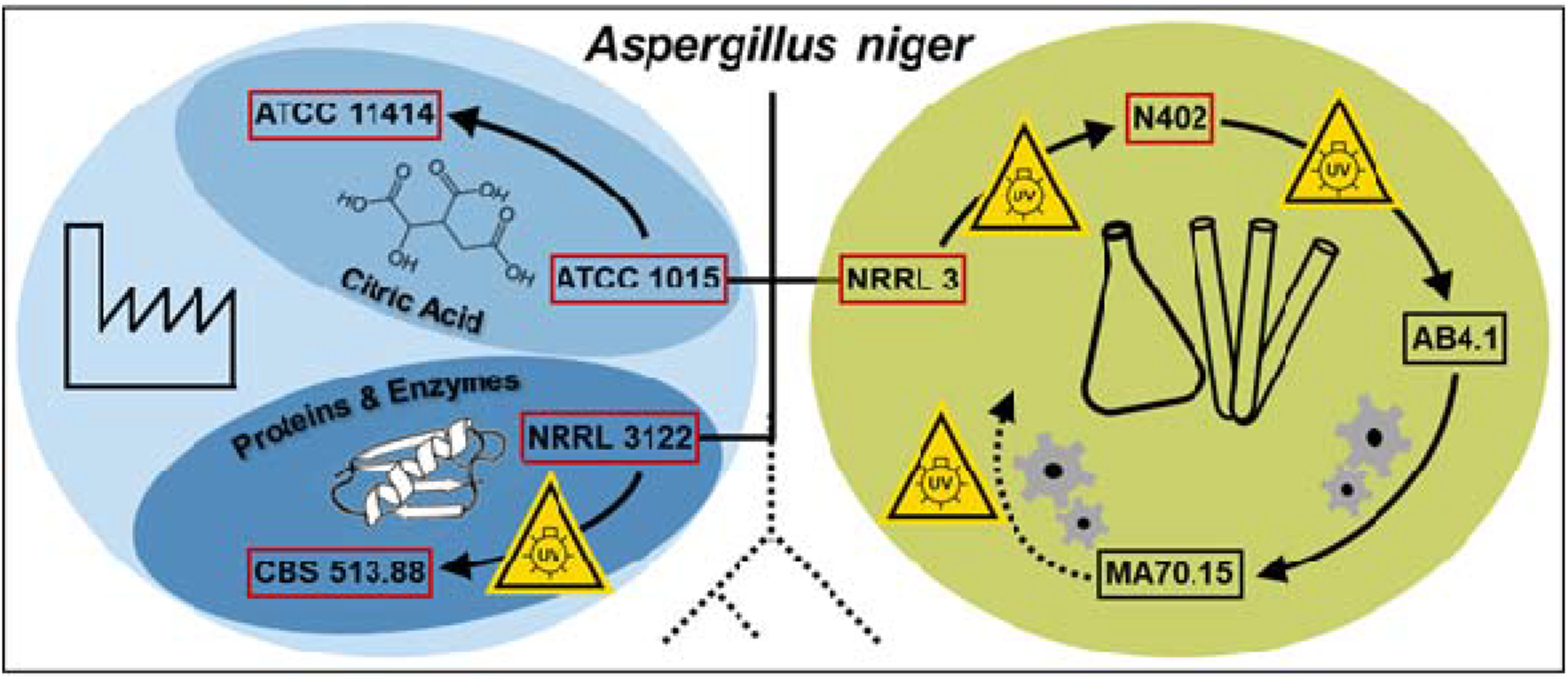
Highly relevant lineages of the *A. niger* species and their primary applications. Schematic overview listing selected but representative members of openly available and related *A. niger* strains based on phylogeny [53]. Depicted are commonly used lab strains (green, right) vs. the industrially adapted strains (blue, left) including the citric acid producer clade (light blue) and the more distantly related enzyme producers (dark blue). Yellow signs indicate instances of UV-mutagenesis, grey gear wheels symbolize targeted genomic engineering steps and red boxes indicate strains used in this study.

Ancestral strains, such as *A. niger* ATCC 1015 or *A. niger* NRRL 3122 (amongst others), have been specifically isolated due to their improved citric acid production characteristics or extracellular glucoamylase activity, respectively, and further optimized via sub-culture isolation [38] or UV-mutagenesis [4,51]. The most commonly used strain for laboratory-based genetic analysis, *A. niger* AB4.1, has also emerged from multiple successive rounds of UV mutagenesis on the original wild type (WT) isolate *A. niger* NRRL 3 [7]. These resulted in a short conidiophore phenotype in strain *A. niger* N402 [8] and generation of the *pyrG* auxotrophic marker [19,50]. However, those mutagenic treatments likely gave rise to additional background mutations influencing gene regulation and impacting diverse phenotypic traits. Additional targeted engineering steps on *A. niger* AB4.1 for improved homologous recombination efficiency via kusA disruption in strain *A. niger* MA70.15 [31] or introduction of auxotrophic markers like HisB, NicB or others [16,34] have further expanded the genetic toolbox in the laboratory. Alteration of the genomic repair machinery or the introduction of auxotrophic markers may however induce stress or alter intracellular regulation and hence divert these strains further from their wild type physiological constitution, as e.g. reviewed for *Saccharomyces cerevisiae* [40].

Genomic sequences are openly available for *A. niger* strains CBS 513.88 [37], ATCC 1015 [4] and NRRL 3 [2] – with recent advances on its annotation [43] – as well as for three additional isolates [53]. Comparison of intra-species genomic data was able to reveal the cause for some of the observed phenotypic differences between the strains, such as overproduction of glucoamylase in *A. niger* CBS 513.88 due to an additional glucoamylase gene acquired by horizontal gene transfer [4,53]. Transcriptional profiling however also indicated that the regulatory networks between different strains are apparently already highly divergent and cannot be explained purely by genomic observations [4]. Furthermore, fungal morphology in submerged cultures was shown to affect fungal productivity for different purposes [10,23,52,55].

Accordingly, none of the current datasets provide enough information to predict superiority of any available strain for the production of pectinases. Moreover, to the best of our knowledge, no thorough comparison for pectinase activity between the available and highly cited *A. niger* strains (listed in Fig. 1) has been conducted so far. While the availability of commercial *A. niger* pectinase cocktails indicates established pectinase production strains in industry, almost no information on their specific origin is publically accessible. In the academic field, numerous studies on optimization of fermentation conditions for pectinase production with various strains have been published (e.g. [1,18,41]). However, these do not allow for direct performance comparison of individual strains due to varying study designs and fermentation conditions. Furthermore, thorough comparisons of different strains under reproducible conditions are scarce in literature and compromised by limited morphology control (due to execution in shake flasks, for example, instead of controlled stirred-tank bioreactors) or poor description of strain origins (e.g. [17,24]).

The importance of pectin, pectinases, and pectin-derived sugars as well as the predominance of *A. niger* in their production processes, however, dictate the necessity for a systematic comparison of strains under controlled and highly reproducible conditions to identify efficient and marker-free host strains for larger scale pectinase production. Therefore, a total of six strains (red boxes in Fig. 1) were selected for comparison of endogenous pectinase activity based on five key prerequisites: (i) relevance to the field based on the number of publications using these strains, (ii) availability of the genomic sequence (or that of a very closely related strain) as a premise for successive genetic optimization, (iii) absence of auxotrophic markers to avoid phenotypic differences due to mutations in central metabolism, (iv) absence of any targeted engineering of elements regulating pectinase expression to avoid distortion of the underlying endogenous pectinase capacity, and (v) classification as biosafety level 1 to allow for universal handling.

By obtaining data on sporulation efficiency, total protein secretion, total and endo-specific polygalacturonase (PGase) activity as well as morphology in submerged culture, this study thus provides essential insights into selection of suitable base strains for pectinase production. Working with controlled stirred-tank bioreactor fermentations after the initial pre-selection, we thereby applied a robust and reproducible methodology rarely employed in phenotypic comparisons of fungal strains, resulting in the identification of a superior *A. niger* strain and potential chassis for additional genetic optimization to boost d-GalA release from complex pectinaceous residues.

## Materials and Methods

### 2.1 Strains, Inoculum preparation and cultivation medium

*A. niger* strains ATCC 1015, ATCC 11414 and NRRL 3122 were obtained from the NRRL collection, NRRL 3, N402 and CBS 513.88 were obtained from the group of Mark Arentshorst at Leiden university. Fungal spores were grown on 39 g L^−1^ potato extract glucose agar (Carl Roth GmbH + Co. KG, Karlsruhe, Germany) supplemented with 10 g L^−1^ yeast extract and 1x trace elements solution, referred to as rich complete medium. After 120 h at 30°C, spores were harvested using sterile 0.89% NaCl solution with 0.05% Tween 80.

All experiments involving submerged fungal cultivation were carried out in 2% (w/w) pectin minimal medium containing (L^−1^): 20.0 g pectin C, 6.0 g NaNO_3_, 1.5 g KH_2_PO_4_, 0.5 g KCl, 0.5 g MgSO_4_ · 7 H_2_O, 1 ml trace element solution and 1 ml PPG P2000 (antifoam, only if cultivated in stirred tank fermenter). The trace element solution was prepared as (L^−1^) 10 g EDTA, 4.4 g ZnSO_4_ · 7 H_2_O, 1.01 g MnCl_2_ · 4 H_2_O, 0.32 g CoCl_2_ · 6 H_2_O, 0.315 g CuSO_4_ · 5 H_2_O, 0.22 g (NH_4_)_6_Mo_7_O_24_ · 4 H_2_O, 1.47 g CaCl_2_ · 2 H_2_O and 1 g FeSO_4_ · 7 H_2_O [5].

### 2.2 Cultivation conditions

#### Shake flask cultivation

The fungi were grown in 250 ml flasks without baffles containing 25 ml of the 20 g L^−1^ pectin minimal medium at 250 min^−1^ (25 mm shaking throw) and 30°C for 96 h. The initial pH was set to pH 4.5 and the cultivation was inoculated to a spore density of 10^9^ spores L^−1^. Strains were grown in triplicates. Data were statistically evaluated by applying an analysis of variance (one-way ANOVA) followed by a Tukey’s post-hoc test using the software Origin (OriginLab). Differences among the mean activity measurements were calculated at a significance level of 0.05 (*p* < 0.05).

#### Bioreactor cultivation

A 7 L stirred-tank bioreactor equipped with three baffles and three six-blade Rushton turbines (Labfors, Infors-HT, Bottmingen, Switzerland) was used during all cultivations. All processes were performed equally under the following conditions. 3 liters of the 20 g L^−1^ pectin mineral medium was inoculated to 10^9^ spores L^−1^. Temperature was kept constant at 30°C. The pH was controlled to a set-point of pH 4.5 by the addition of either 1M H_2_SO_4_ or 3M KOH. Batch processes were carried out for 86 h to 90 h. To prevent initial spore loss, the stirred-tank bioreactor was not aerated and only slowly mixed at 250 min^−1^ (~0.13 W L^−1^ [20]) during the first 6 h of batch cultivations [33]. Afterwards, the stirrer speed was set to 700 min^−1^ (~1.625 W L^−1^ [20]) and aeration to 0.2 vvm, which was also sufficient to keep the dissolved oxygen concentration above 30% air saturation during all cultivations conducted. Additionally, exit-gas composition (O_2_, CO_2_) was monitored (EasyLine, ABB, Zürich, Switzerland).

### 2.3 Biomass dry weight concentration

Biomass dry weight was determined by filtering a known volume thought pre-dried and pre-weighted filter paper (Whatman No. 1 & 5). The collected biomass was dried at 90°C to constant weight and reweighted. The determination of the biomass dry weight was performed in triplicate and pictured as mean with standard deviation of the measurements.

### 2.4 Morphological characterization

Microscopic images for morphological characterization were taken with an Axioplan microscope (Carl Zeiss AG, Jena, Germany) at 1.25x magnification directly after sampling after 9 h, 12 h, 19 h, 36 h and 88 h of the batch cultivation. The microscope was equipped with a 3.3-megapixel Axiocam ICc3 microscopy camera (Carl Zeiss AG, Jena, Germany).

### 2.5 Protein concentration of culture supernatant

Protein concentration of culture supernatant was determined by Coomassie (Bradford) Protein Assay Kit (Thermo Scientific) according to manufacturer’s specifications. Each sample was diluted with 0.1 M sodium citrate buffer pH 4.5, mixed with the Bradford reagent and incubated for at least 10 min at room temperature. Afterwards, the absorbance at 595 nm was measured with a multimode microplate reader (Infinite M200, Tecan, Männedorf, Germany). Bovine serum albumin was used as standard. The determination of the protein concentrations was performed in triplicate and pictured as mean with standard deviation of the measurements.

### 2.6 Total polygalacturonase activity

The pectinase activity was determined following a miniaturized version of the Fructan Assay Kit protocol (Megazyme) for reducing sugars. 10 μL of the culture supernatant and 10 μL of a 5 g L^−1^ polygalacturonic acid solution (PGA, buffered in 0.1 M sodium citrate, pH 4.5) were mixed and incubated for 40 min at 30°C. The released reducing sugar ends were determined using a 4-hydroxybenzhydrazide solution as described in the Megazyme protocol and measured at 410 nm with a multimode microplate reader (Infinite M200, Tecan, Männedorf, Germany).Sample values were blanked against similarly prepared but non-incubated mock samples. One unit of total polygalacturonase activity equals the amount of enzyme that catalyzes the formation of one μmol of d-galacturonic acid per minute under assay conditions. Assaying was performed in triplicate and plotted as means with standard deviation of replicate measurements.

### 2.7 Endo-polygalacturonase activity

Endo-polygalacturonase activity was assessed following the protocol of Ortiz [35] 8 μl of a 5 g L^−1^ polygalacturonic acid solution (PGA, buffered in 0.1 M sodium acetate, pH 4.5) and 8 μl of *A. niger* culture supernatant were mixed and incubated for 30 min at 30 °C in a microtiter plate prior to adding 40 μl of freshly prepared ruthenium red working solution (Sigma Aldrich, 1.125 mg mL^−1^ in ddH_2_O) and 100 μl of 8 mM sodium hydroxide solution. Samples were spun down at 3200 g for 10 min and diluted in a one to eight ratio for absorbance measurement of 200 μl diluted sample at 535 nm on a microplate reader (Infinite M200, Tecan, Männedorf, Germany). Sample values were blanked against similarly prepared but non-incubated mock samples. One enzyme unit was defined as the amount of enzyme required to hydrolyze 1 μg of polygalacturonic acid in smaller fragments unable to precipitate with the dye per minute under the assay conditions. Assaying was performed in triplicate and plotted as means with standard deviation of replicate measurements.

### 2.8 Hydrolysis of sugar beet pulp

The hydrolysis of pectin-rich residues was conducted with pre-dried (50°C) and milled sugar beet press pulp provided by Südzucker AG (Mannheim, Germany). 10 g of pre-sterilized pulp was mixed with 100 ml of reaction solution. The reaction solution consisted of 90 ml of sterile filtered (0.22 μm) enzyme supernatant of the strains *A. niger* N402 (172.48 ± 3.90 mg L^−1^ Protein; 206.62 ± 26.00 U L^−1^ total PGase activity) or *A. niger* ATCC 11414 (111.26 ± 9.60 mg L^−1^ Protein; 1475.69 ± 32.28 U L^− 1^ total PGase activity) buffered with 10 ml of 1.0 M sodium acetate (pH 4.5). Hydrolysis was carried out at 180 rpm min^−1^ and 30°C for 138 h in sterile and closed 250 ml glass bottles (DWK Life Sciences GmbH, Mainz, Germany) to prevent evaporation and contamination. Time series samples were taken from homogenized hydrolysis mixes and stored at −80°C until further use.

### 2.9 HPAEC-PAD analysis of the hydrolysis supernatant

Free d-GalA amounts in hydrolysis samples (diluted 1:4000) were determined on a Dionex ICS 3000 HPAEC-PAD instrument setup with Dionex AS Autosampler, Dionex gradient mixer GM-3 (Dionex Corp., Sunnyvale, California, USA) and CarboPac PA1 standard bore guard column (4 x 50 mm) plus CarboPac PA1 preparative IC column (4 x 250 mm, both Thermo Fisher Scientific Inc., Waltham, Massachusetts, USA) using a 12.5 min linear gradient of 100 to 250 mM sodium acetate in 100 mM sodium hydroxide solution (prepared in low total organic carbion deionized water) at 1 ml min^−1^ flow rate and constant 30 °C elution temperature.

## Results and Discussion

### 3.1 Shake flask-based pre-selection of *A. niger* strains for pectinase production

As a first step in the identification of an ideal base strain for pectinase production among the selected *A. niger* strains, an initial fast and cost-efficient experiment was conducted in small-scale using shake flasks as typically applied in strain screenings (e.g. [39]). Total polygalacturonase (PGase) activity was measured in supernatants of *A. niger* cultures after 96 h of cultivation in minimal medium supplemented with 2% (w/v) pectin C as a carbon source (Fig. 2A). Strains *A. niger* NRRL 3122 and *A. niger* CBS 513.88 repeatedly revealed the lowest total PGase activities of all strains tested (174.43 ± 41.22 U L^−1^ and 344.96 ± 199.34 U L^−1^, respectively), while *A. niger* NRRL 3 (606.95 ± 173.88 U L^−1^), *A. niger* N402 (493.17 ± 233.44 U L^−1^), *A. niger* ATCC 1015 (674.36 ± 92.38 U L^−1^) and *A. niger* ATCC 11414 (543.18 ± 245.95 U L^−1^) revealed superior total PGase activities. Additionally, *A. niger* NRRL 3122 and *A. niger* CBS 513.88 displayed inferior spore densities on rich medium, which was rated as a disadvantage for larger scale liquid culture inoculations (Fig. 2B). Based on the combination of these results, strains *A. niger* NRRL 3122 and *A. niger* CBS 513.88 were excluded from further tests.

**Fig. 2:**
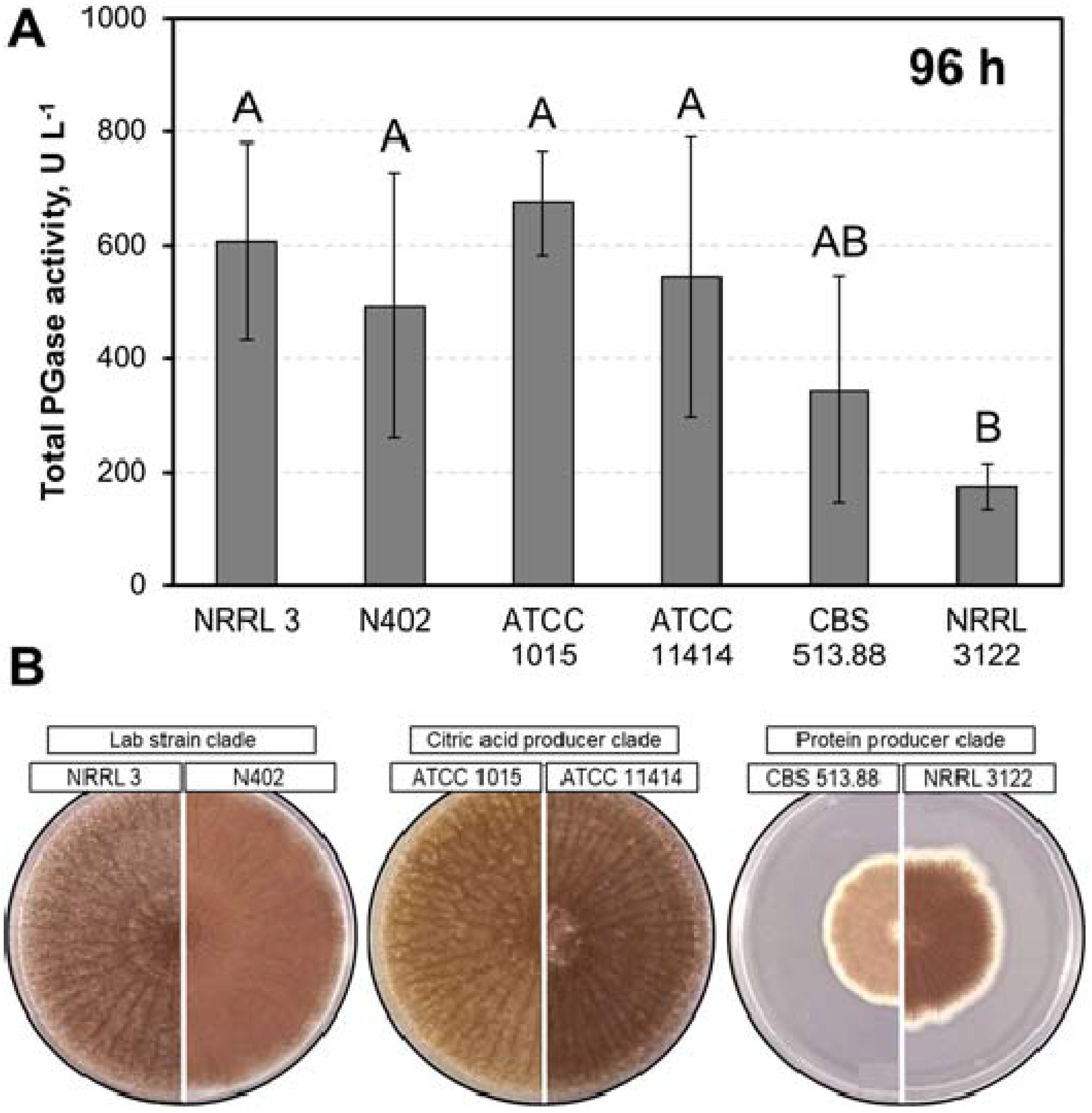
Polygalacturonase activity and sporulation density of *A. niger* screening strains. (A) Total PGase activity of *A. niger* culture supernatants of a representative shake flask batch after 96 h of incubation in minimal medium + 2% (w/v) pectin C. Different capital letters indicate significant differences within the displayed data groups (*p* < 0.05) using a one-way ANOVA followed by a Tukey’s post-hoc test. Group AB showed no significant differences either to group A and group B. (B) Sporulation densities of all strains on rich complete medium after six days.

*A. niger* CBS 513.88 (and its ancestor *A. niger* NRRL 3122) particularly differ from *A. niger* ATCC 1015 and its closely related *A. niger* NRRL 3 lineages by additionally acquired glucoamylase (*glaA*) genes from horizontal gene transfer as well as the upregulation of amino acid synthases that are overrepresented in GlaA and their respective tRNAs [4]. Considering their poor performance in this study however, *A. niger* CBS 513.88 and *A. niger* NRRL 3122 do not seem to have a universal advantage over the other tested strains in terms of protein production and secretion *per se*, but rather a limited one for GlaA expression.

### 3.2 Comparison of pre-selected *A. niger* strains in submerged stirred-tank bioreactor batch cultivations

A high degree of control over mechanical and physicochemical parameters influencing submerged culture morphology as well as enzyme activity were shown to be important for achieving high pectinase activity [1,14,52]. In a next step, pectinase production of all strains was therefore evaluated in batch processes on a 3 L scale in controlled stirred-tank bioreactor cultivations to perform selection under robust and reproducible conditions (see Figs. 3, 4). To this end, submerged batch cultivations in 2% (w/v) pectin minimal medium were conducted to investigate the pectinolytic properties of the four best performing *A. niger* strains from the pre-selection. For each batch cultivation, biomass dry weight concentration (BDW), total protein concentration (c_Protein_), total polygalacturonase and endo-polygalacturonase (PGase) activity as well as fungal morphology were determined, all being highly relevant variables for strain productivity in submerged cultures [28,52]. Since the level of morphology control is generally higher in stirred tank bioreactors compared to shake flasks, we see our bioreactor approach as advantageous for in-depth strain comparison after pre-selection [14].

**Fig. 3:**
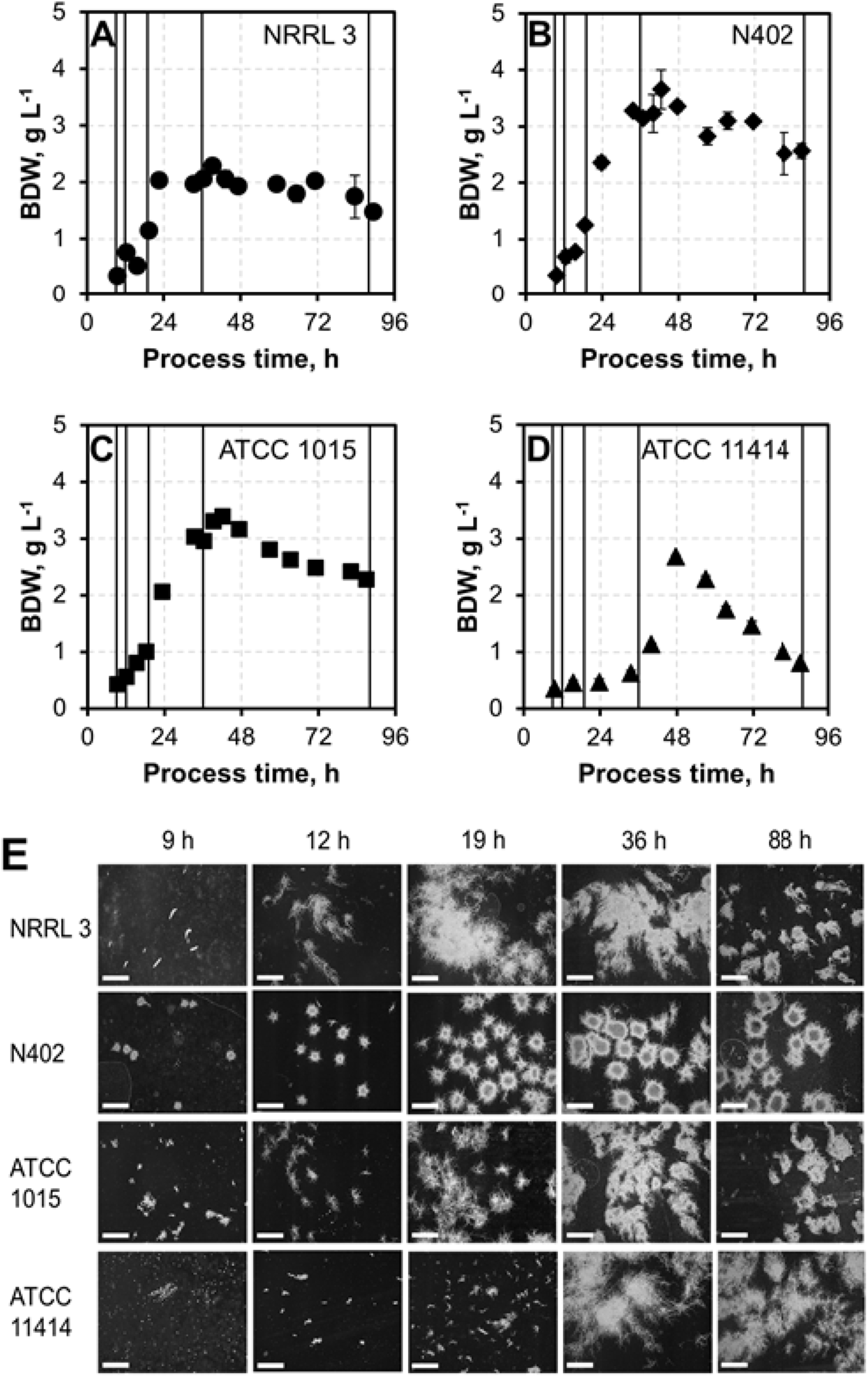
Biomass dry weight concentrations (BDW) and morphology of selected *A. niger* strains. (A – D) BDW of *A. niger* NRRL 3 (A, •), *A. niger* N402 (B, ♦), *A. niger* ATCC 1015 (C, ■) and *A. niger* ATCC 11414 (D, ▲) during 90 h submerged batch cultivations in a 3 L stirred-tank bioreactor with 2% pectin minimal medium. (E) Morphological changes of all four *A. niger* strains throughout the cultivations are shown below. White scale bars indicate 1 mm. Lines in A – D indicate morphology sampling times.

**Fig. 4:**
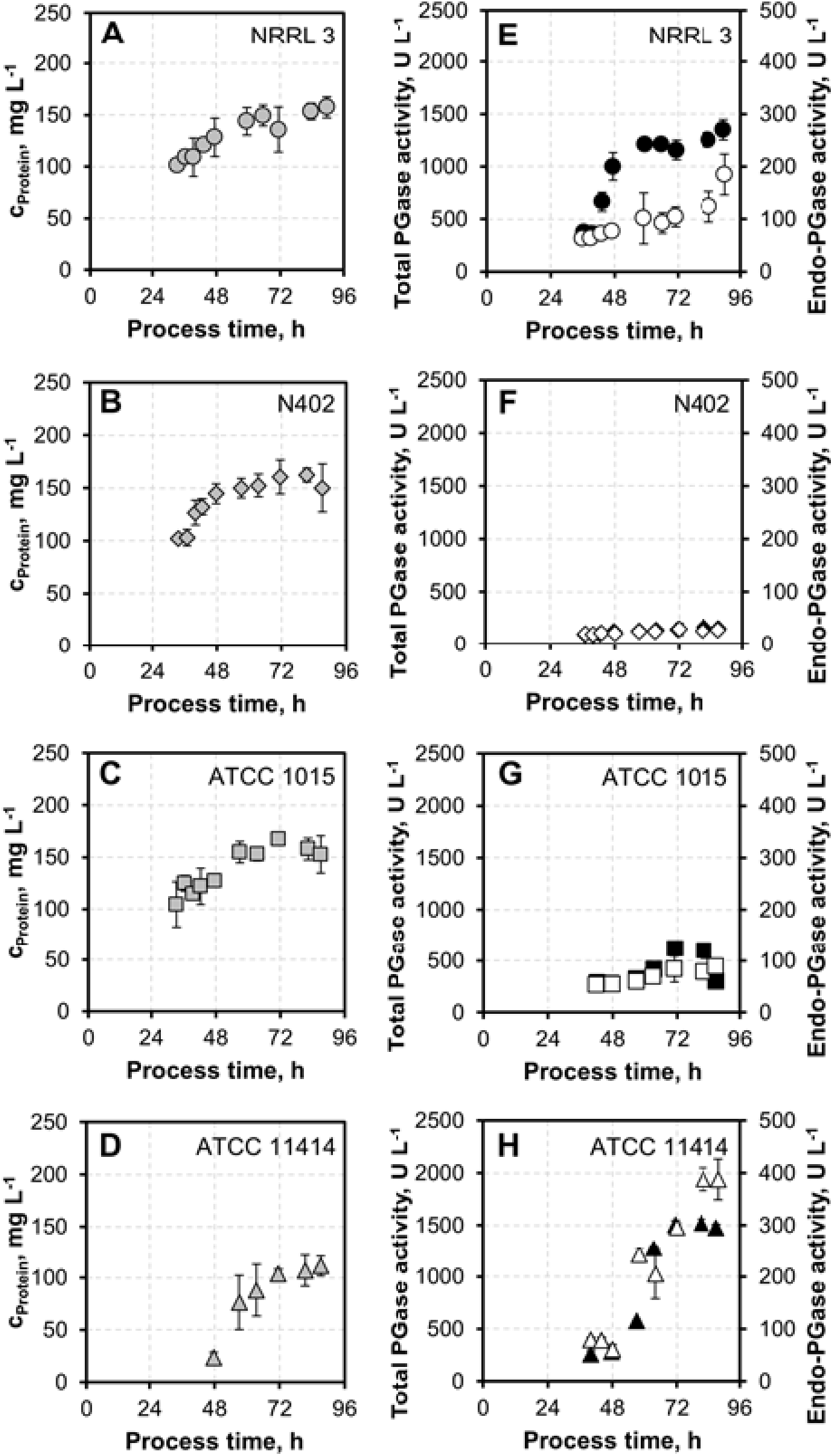
Protein concentrations, total and endo-PGase activities of pre-selected *A. niger* strains in 3 L controlled stirred-tank batches. (A – D) Total secreted protein concentrations (grey) in culture su-pernatants of pre-selected *A. niger* strains in 2% pectin minimal medium over a time course of 90 h. (E – H) Total (black) and endo-PGase activities (white) of culture supernatants throughout cultivation.

### 3.2.1 Biomass dry weight concentrations and morphology

In case of *A. niger* NRRL 3, BDW rose to its maximum at 2.28 ± 0.12 g L^−1^ during the first 39 h and slightly decreased about 35% afterwards until the end of the cultivation (1.47 ± 0.01 g L^−1^) (Fig. 3A). The BDW of the strains *A. niger* N402 and *A. niger* ATCC 1015 showed a similar behavior, increasing to 3.65 ± 0.35 g L^−1^ (*A. niger* N402) and 3.38 ± 0.07 g L^−1^ (*A. niger* ATCC 1015) within 42 h before decreasing by 30% until the end of the cultivation (Fig. 3B,C). A different behavior could be observed for strain *A. niger* ATCC 11414. Its BDW peaked only after 48 h (at 2.68 ± 0.03 g L^−1^), with a drastic decrease of 70% to 0.81 ± 0.02 g L^−1^ at the end of the observation period (Fig. 3D). The loss of BDW in the cultivations is in accordance with the dissolved oxygen concentration (DO) and the carbon dioxide fraction in the exit gas measured online. DO increased with initiating BDW loss and the carbon dioxide fraction in the exit gas decreased without change in the aeration of the process, indicating a limitation of the energy source (data not shown).

Since the productivity of filamentous fungi in submerged cultivations was shown to be highly dependent on their morphology [13,23,36,55], monitoring of this parameter was conducted throughout each submerged cultivation. Fig. 3E depicts the morphology of all four strains after 9 h, 12 h, 19 h, 36 h and 88 h of the cultivation. After 36 h, a stable morphology was observed for all strains. *A. niger* NRRL 3 and *A. niger* ATCC 11414 showed disperse, mycelial-like growth, *A. niger* N402 strongly pellet-like growth and *A. niger* ATCC 1015 a less dense but still pelleted form of growth. Towards the end of cultivation, the morphology of *A. niger* NRRL 3 shifted to a more pelleted structure. Notably, strong yellow pigmentation matching the description of well-known Aurasperone formation by *A. niger* [46] occurred after 34 h in all culture supernatants with the exception of *A. niger* ATCC 11414, where only mild pigmentation was observed after 58 h. As all four batch cultivations were run under identical conditions, these results indicate severe intra-species differences in physiological regulation between the selected strains that cannot be explained by comparison of their genomic sequences alone [4].

### 3.2.2 Secreted protein concentrations and PGase activities

Next, the total secreted protein concentration in the supernatants (Fig. 4A-D) as well as the total and endo-specific PGase activities were assessed (Fig. 4E-H). The protein concentration of the cultivations with *A. niger* NRRL 3, *A. niger* N402 and *A. niger* ATCC 1015 displayed very similar development with a mildly logarithmic-like increase throughout the cultivation, peaking at 157.39 ± 10.22 mg L^−1^ (90 h), 162.34 ± 6.38 mg L^−1^ (81 h) and at 167.43 ± 5.35 mg L^−1^ (71 h), respectively. Following the observed BDW decrease towards the end of cultivation, *A. niger* N402 and *A. niger* ATCC 1015 also displayed a mild decrease in protein concentration. In accordance with its delayed biomass generation (Fig. 3D), *A. niger* ATCC 11414 showed its maximal protein titer (111.26 ± 9.60 mg L^−1^) only after 87 h (Fig. 4D). This protein concentration was 32% lower than the maximal concentration observed for the other three investigated strains. In contrast to predominant observations for a variety of secreted proteins expressed in *A. niger* and other filamentous fungi [28,52], dispersed mycelial growth therefore did not prove to be a direct indicator for increased overall protein secretion across different strain lineages of *A. niger*. However, considering specific secretion rates (normalized to fungal biomass), *A. niger* ATCC 11414 is performing *on par* with the other strains – particularly towards the end of the incubation time.

Intriguingly, in terms of PGase activities, the dispersely growing *A. niger* NRRL 3 and *A. niger* ATCC 11414 strains showed superior performance. Total PGase activity of *A. niger* NRRL 3 showed a sharp increase between 36 h and 59 h and a maximum activity of 1378.87 ± 98.39 U L^−1^ (90 h). Endo-PGase activity of *A. niger* NRRL 3, however, showed a rather exponential increase to a maximum of 184.55 ± 38.82 U L^− 1^ throughout the cultivation and appeared to be decoupled from the total PGase activity, indicating differences in expressional regulation of individual pectinase classes. The highest total PGase activity was generated by *A. niger* ATCC 11414, continuously increasing throughout the cultivation to a maximum of 1524.17 ± 34.90 U L^−1^ (82 h). Endo-PGase activity peaked at 388.61 ± 21.64 U L^−1^ at 82 h. *A. niger* ATCC 11414 thus exceeded the maximal total PGase activity of NRRL 3 – as the second strongest polygalacturonase producer - by 13% and the maximal endo-PGase activity by 111%. Total secreted protein as well as biomass generation hence could not be used as a proxy for pectinase activity in this study. Moreover, the different maximum activities for endo- and total PGase as well as the different time profiles of PGase activity for *A. niger* NRRL3 and *A. niger* ATCC 11414 throughout the cultivation indicate severe differences in the regulation and expression of pectinases between these two strains. Considering the lower biomass accumulations and total protein production during the cultivation of *A. niger* ATCC 11414, this strain moreover generated the highest specific total and endo-PGase activities compared to all of the other investigated strains.

Assuming similar total capacities of the secretory machineries in all tested *A. niger* strains, *A. niger* ATCC 11414 is therefore recognized as the most promising candidate for genetic improvements towards pectinase overexpression. Less off-target secondary metabolism activity (as judged by pigment formation) and highest specific polygalacturonase production hold promise for additional metabolic capacities which might be exploitable for enhanced pectinase expression.

## 3.3 Comparison of hydrolytic performance of culture supernatants for d-GalA release from complex pectinaceous substrates

Based on the results presented above, *A. niger* ATCC 11414 was determined as the most promising base strain for pectinase production out of six strains under the tested cultivation conditions. *A. niger* ATCC 11414 not only had the highest total and specific PGase activity, but also showed a disperse morphology desirable for protein secretion.

To test whether the selection of *A. niger* ATCC 11414 based on defined substrate assay conditions would translate into improved activity also on complex pectinaceous biomass, d-GalA release from milled dry sugar beet press pulp (SBPP) using *A. niger* ATCC 11414 culture supernatant was compared against *A. niger* N402 culture supernatant as the ancestor of today’s standard laboratory strains. Using a 96 h *A. niger* ATCC 11414 buffered culture, an average of 8.8 g L^−1^ of free d-GalA was released from 9% (w/v) SBPP within 138 h, as compared to an average of 4.9 g L^−1^ for *A. niger* N402 culture supernatant (Fig. 5). Taking into account the water molecules incorporated during hydrolytic cleavage, this corresponded to a degraded amount of 8.0 g L^−1^ polygalacturonan and 4.5 g L^−^1 of the provided biomass, respectively. Considering a total d-GalA content in SBPP of approximately 22% (w/w) ([26]), ~36.4% of the expectable d-GalA could be released using *A. niger* ATCC 11414 culture supernatant (vs. ~20% with *A. niger* N402 culture supernatant). In other words, the same release level was realized in less than 45% of the process time. In summary, thorough screening and activity-driven selection of *A. niger* strains from a set of readily available and highly referenced strains resulted in a 75% higher d-GalA release compared to a scenario in which the standard lab strain was used.

**Fig. 5:**
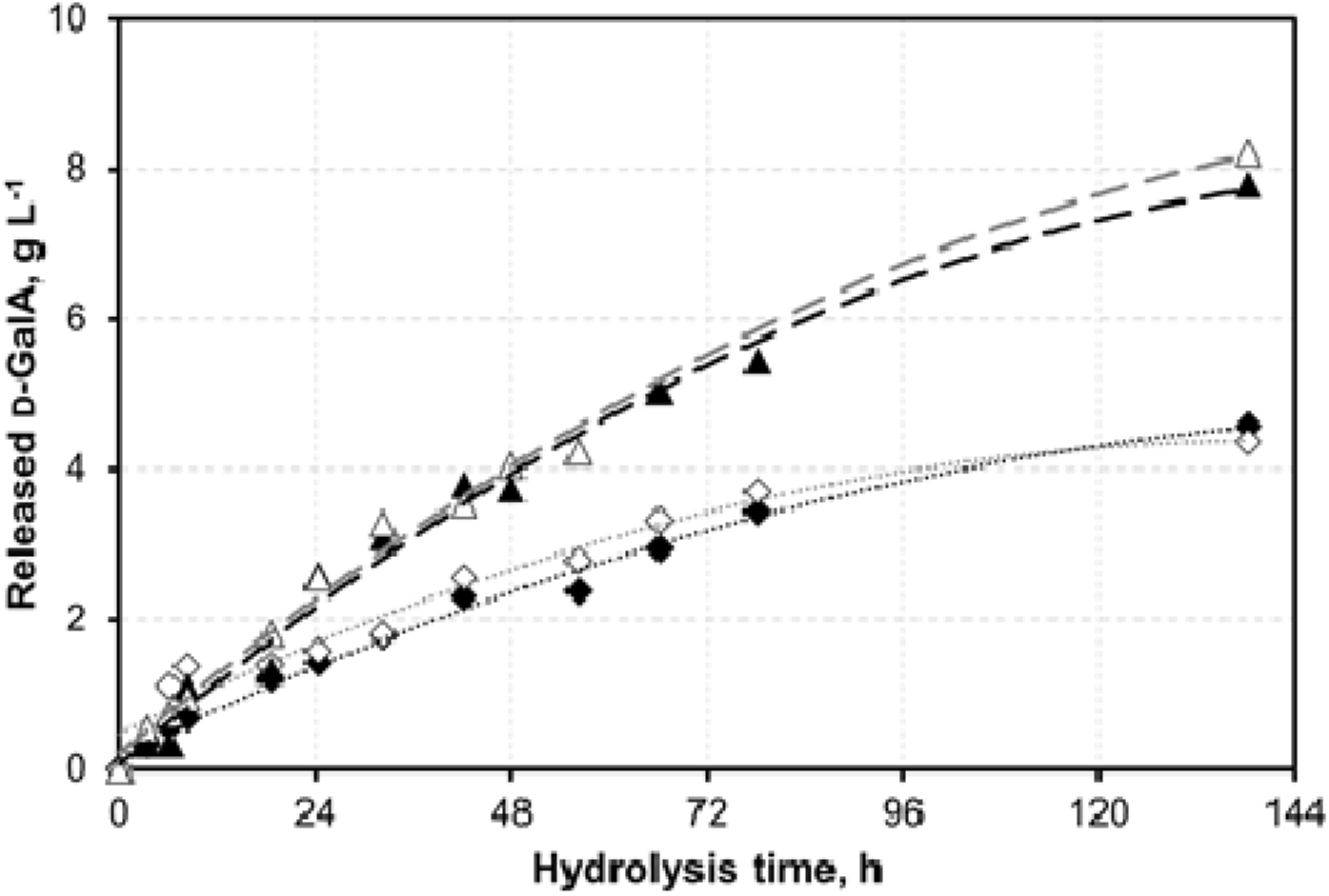
d-GalA release from 9% sugar beet press pulp (SBPP) using *A. niger* ATCC 11414 vs. *A. niger* N402 culture supernatant. Black and white triangles (Δ, ▲) represent replicates of hydrolysis using 96 h culture supernatants of *A. niger* ATCC 11414 stirred tank batch cultivations in 2% pectin minimal medium, with respective dashed lines in grey and black. Black and white diamonds (♦, ◊) represent two replicates using supernatants of N402 with respective dotted lines in black and grey.

Monomeric d-GalA release from SBPP of up to 79% within 48 h has been reported in highly optimized saccharification conditions using combinations of commercial pectinase mixes of the *Aspergillus* genus at comparable enzyme concentrations as used in this study [29], while no comparable efficiency data exist for direct application of crude *Aspergillus* culture supernatants on SBPP. Furthermore, the hydrolysis setup presented in this study was used as a pectinase production benchmark in strain selection only and has not yet undergone optimization for ideal d-GalA release conditions. Additional optimization of culture conditions for *A. niger* ATCC 11414 may further improve its performance, *e.g.* in terms of protein secretion, for which titers of up to 20 g L^−1^ have been reported [32]. Supplementation of (hemi-)cellulosic substrates and thereby induced expression of (hemi-)cellulases during pectinase production could further contribute to d-GalA release from complex biomasses [11,29]. Research in this field is still actively ongoing, but mostly focusing on process engineering, as recently demonstrated for continuous generation of d-GalA from SBPP pectin extracts in membrane enzyme reactors [15]. Via systematic screening and strain performance evaluation under controlled conditions, we complemented this research with an important comparison of highly cited, openly available and readily applied *A. niger* strains. Endeavors to use SBPP and other complex pectinaceous biomasses of interest for industrial d-GalA supply in the context of the bioeconomy hence could highly benefit from this work.

## Conclusion

Considering the lack of systematic screening for strong pectinase producing *A. niger* strains, we have implemented a robust protocol for the discrimination of competing strains in controlled stirred-tank bioreactors. Superior performance of *A. niger* ATCC 11414 was verified in a realistic setting using complex sugar beet press pulp. This strain shows potentially untapped metabolic and secretory reservoirs that could be exploited for improved pectinase production via targeted genetic engineering. However, to foster transfer of research results to industrial applications, it will be necessary to establish genetic tools, such as non-homologous end-joining suppressors or genetic markers, in this non-standard host strain.

E-supplementary data of this work can be found in online version of the paper.

## Acknowledgements

This work was supported by the German Federal Ministry of Education and Research [grant numbers 031B0342A and 031B0342C]. We furthermore want to thank Petra Arnold, Sabrina Paulus and Nadine Griesbacher (Wood Research Munich, TUM, Germany), as well as Markus Amann und Norbert Werth (Institute of Biochemical Engineering, TUM, Germany) for excellent technical assistance. Also, we would like to acknowledge Nathalie Hafner (Institute of Biochemical Engineering, TUM, Germany) for assisting with the assays and fermentations. Additionally, we want to acknowledge the Südzucker AG, Mannheim, Germany for kindly providing the sugar beet press pulp and Mark Arentshorst (Leiden University, Netherlands), Vera Meyer and Tabea Schütze (both TU Berlin, Germany) for providing strains and lab trainings for this study. The support of Dominik Schäfer and Kevin Schmitz by the TUM Graduate School is acknowledged as well.

## Supplementary Figures

**Fig. S1:**
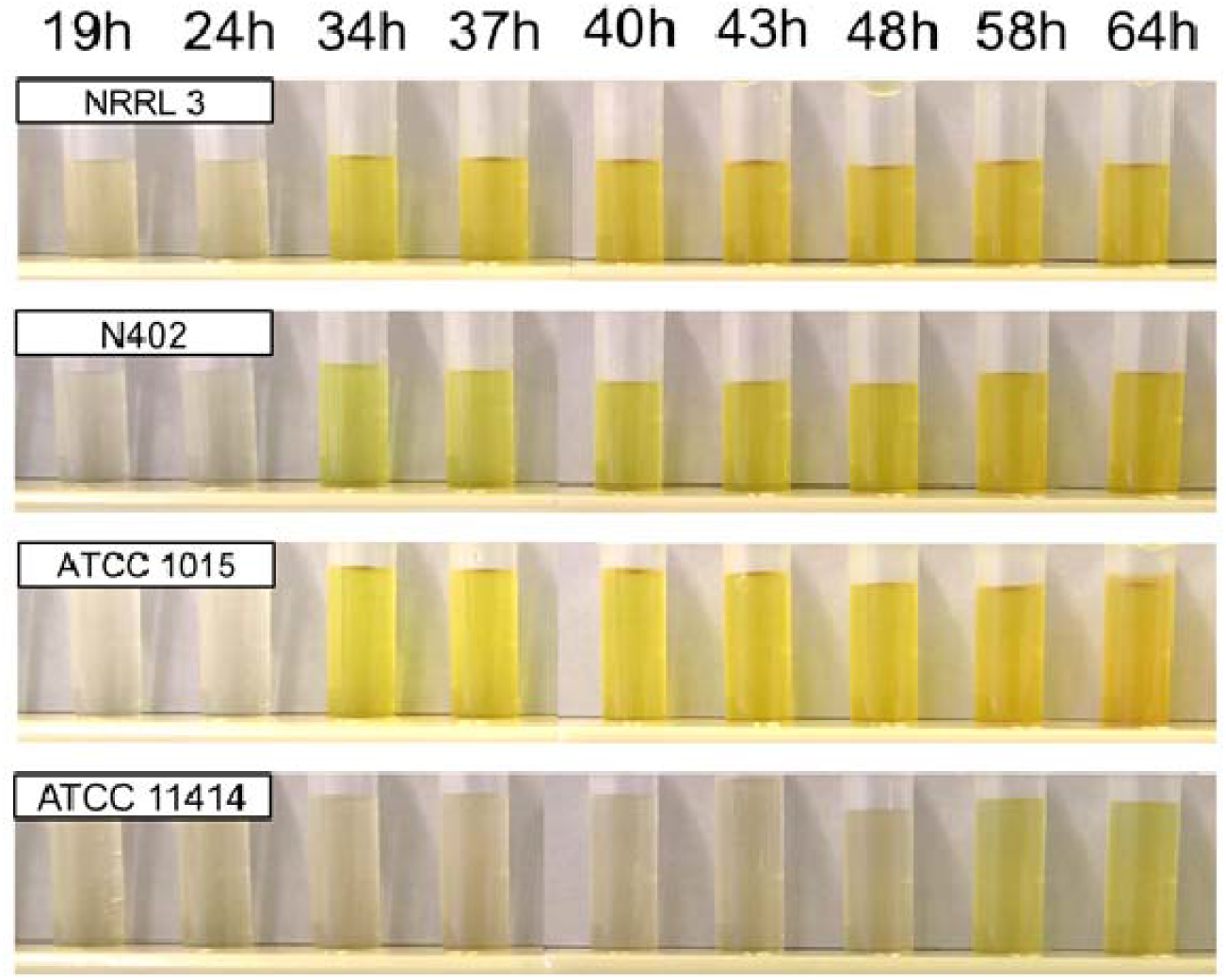
Yellow pigment formation in *A. niger* culture supernatants. Accumulation of yellow pigmentation in culture supernatants of *A. niger* NRRL3, N402, ATCC1015 and ATCC11414 submerged stirred tank bioreactor batch cultivations in 2% pectin minimal medium.

**Fig. S2:**
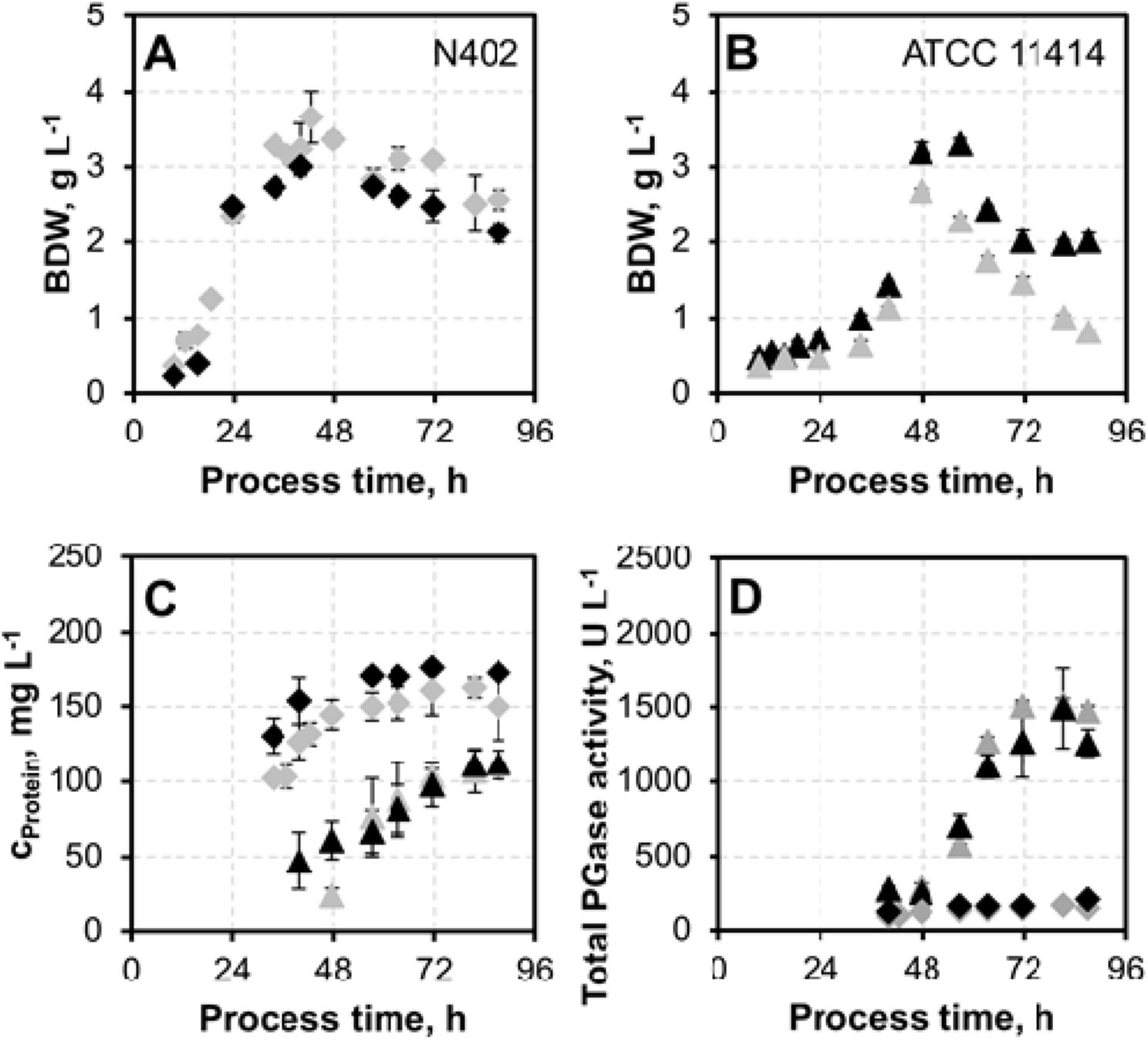
Reproduced submerged stirred tank batch cultivations of *A. niger* N402 and *A. niger* ATCC 11414. Biomass dry weight concentrations (A, B), total protein concentration (C) and total PGase activity (D) of the reproduced cultivations of *A. niger* N402 (◆) and *A. niger* ATCC 11414 (▲). Grey symbols display the values of the cultivations depicted in the results and discussion.

**Fig. S3:**
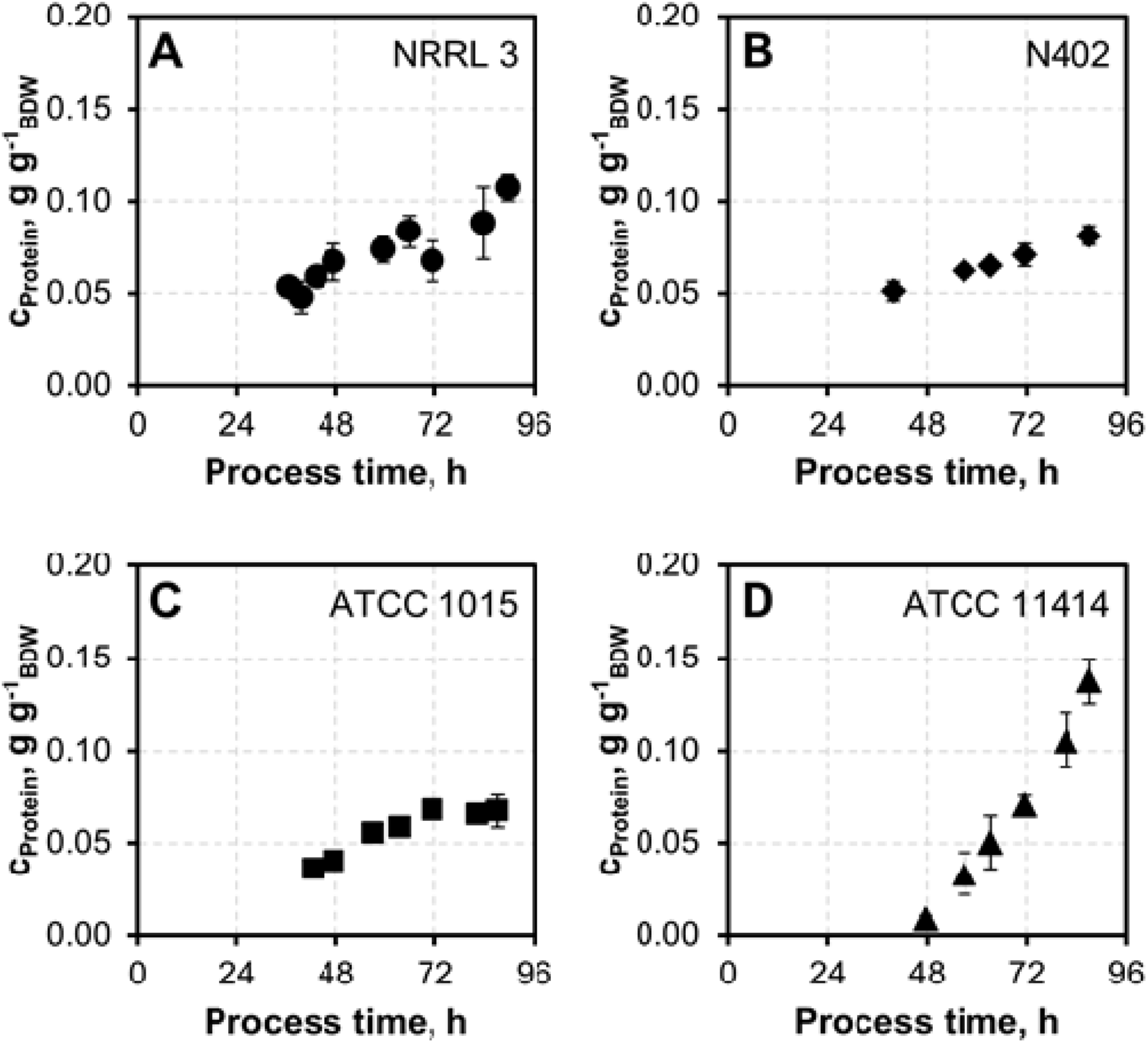
Specific protein concentrations of selected *A. niger* strains. *A. niger* NRRL 3 (A, ●), *A. niger* N402 (B, ◆), *A. niger* ATCC 1015 (C, ■) and *A. niger* ATCC 11414 (D, ▲) specific protein concentrations normalized by BDW during 90 h submerged batch cultivations in a 3 L stirred tank bioreactor in 2% pectin minimal medium.

